# Poultry-Associated Nitrofurantoin-Resistant and Pre-Resistant *Escherichia coli* Clones are Found in Multiple Countries and One-Health Compartments

**DOI:** 10.1101/2024.10.20.619276

**Authors:** Jordan E Sealey, Beth Astley, Oliver Mounsey, Matthew B Avison

**Affiliations:** University of Bristol School of Cellular & Molecular Medicine, Biomedical Sciences Building, University Walk, Bristol BS8 1TD, United Kingdom

**Keywords:** Zoonosis, Genomics, Urinary tract infection

## Abstract

*Escherichia coli* is the most common cause of urinary tract infection (UTI) in humans. The nitrofuran-class antibacterial drug nitrofurantoin is a frequent UTI therapy, with resistance rarely observed. Here we show that nitrofurantoin resistant (NFT-R) *E. coli* are sometimes excreted by dogs fed a raw meat diet in the city of Bristol, United Kingdom, and that NFT-R and pre-resistant (one mutation away from NFT-R) *E. coli* can be found contaminating chicken meat sold for human consumption and chicken-based raw dog food in the same city. Using whole genome sequencing, we identified multiple NFT-R or pre-resistant *E. coli* clones spanning several phylogroups. These clones were dominated by isolates from poultry farms and poultry meat in Europe, Canada, the United States and Japan, and we identified instances where closely related NFT-R and pre-resistant isolates have colonised humans and caused UTIs. The origins of these poultry-associated NFT-R and pre-resistant *E. coli* clones are uncertain, but nitrofuran-class antibacterials (particularly furazolidone, furaltadone, and nitrofurazone) were used in poultry production during the 1970s and 80s, though this practice has been banned since the 1990s. It is possible, therefore, that this caused an initial selective pressure for the emergence of NFT-R and pre-resistant *E. coli* clones on poultry farms. Our findings have potentially important implications for domestic hygiene, particularly among people receiving nitrofurantoin therapy.

## 1. Introduction

Nitrofurantoin is a nitrofuran-class antibacterial recommended as first line treatment for uncomplicated urinary tract infection (UTI) in adults in the UK, most of which are caused by *Escherichia coli* [1]. Despite widespread use, nitrofurantoin resistance is rare [2].

The ability of nitrofurantoin to kill relies on either of two oxygen-insensitive nitroreductases produced in the bacterium, NfsA and NfsB, which reduce nitrofurantoin to liberate highly reactive intermediates [3]. Nitrofurantoin resistance occurs predominately by loss of function mutations, deletions or insertional inactivation of both *nfsA* and *nfsB*, and in rare cases, by mutation of *ribE* or carriage of the plasmid-mediated *oqxAB* efflux pump genes [2, 4-6]. We have recently identified the presence of nitrofurantoin “ pre-resistant” urinary *E. coli*, where one gene (more commonly *nfsA*) is disrupted, but the other remains intact. The nitrofurantoin MIC increases, but not above the breakpoint defining clinical resistance [2].

The requirement for mutations in two genes – *nfsA* and *nfsB* – may form a barrier to nitrofurantoin resistance emerging among *E. coli* in an infection [7] and there is currently no evidence for nitrofurantoin-resistant (NFT-R) *E. coli* clones circulating in the human population [2]. If such clones emerge, this could spark a rise in NFT-R UTIs.

Nitrofuran-class antibacterials (especially furazolidone and furaltadone, though nitrofurantoin itself was never licenced) were used in food producing animals in Europe, particularly poultry, until the 1990s when the European Union banned this due to the risks associated with toxic chemical residues in meat. Similar bans are in place in the USA, Australia, the Philippines, Thailand and Brazil [8]. In some countries, nitrofurantoin and other nitrofuran-class antibacterials are used to treat ornamental birds, homing pigeons and fish, and as an off-label therapy for recurrent UTI in dogs, cats and horses [9], however, in the UK, nitrofurantoin is not licensed for use in any animal [10].

Surveys of NFT-R bacteria from animals are scarce, and in the few studies that have considered NFT-R *E. coli*, resistance was very rare or non-existent [11-14], but emergence of NFT-R *E. coli* in dogs during treatment of UTI has been reported [15].

We have established a 50 x 50 km study region in the South West of England in which we have sequenced resistant *E. coli* from human infections [2,16-18], and from the faeces of farmed [17,19], companion [20,21], and zoo animals [22]. In the work reported here, we investigated whether NFT-R *E. coli* are excreted by dogs in our study region or are found on meat being sold in stores in the area, and used whole genome sequence (WGS) data to test whether these NFT-R *E. coli* are shared across One Health compartments, including human infections. In doing so, we have identified a reservoir of NFT-R and pre-resistant *E. coli* clones from numerous sequence types (STs) on poultry meat, poultry farms and in dogs fed raw meat. Members of these poultry-associated clones have caused human UTIs.

## 2. Materials and Methods

### 2.1. Canine faecal sample collection and processing

Ethical approval was obtained from the Faculty of Health Research Ethics Committee, University of Bristol (ref: 89282) alongside Ethical Approval of an Investigation Involving Animals (ref: UB/19/057). Details of canine faecal sample collection are published [20,21], but briefly, 297 dogs were recruited at dog-walking sites in the city of Bristol between September 2019 and September 2020. All owners gave informed consent to participate. Faecal samples were collected and a standardized questionnaire was provided to collect demographic data and variables chosen as being potentially associated with carriage of ABR *E. coli* in dogs. Samples were refrigerated and processed within 48 h. Faecal sample aliquots were weighed (0.1– 0.5 g) and PBS was added at 10 mL/g before vortexing and mixing with an equal volume of 50% sterile glycerol. Samples were stored at -70°C.

### 2.2. Meat sample collection and processing

Details of meat sample collection and processing are published [23], but briefly, we sampled beef, chicken, lamb and pork meat collected from large chain grocery stores in the city of Bristol between October and November 2022, totalling 15 samples per type of meat, except for pork where due to availability only 13 were collected. Samples were either processed on the day or kept at 4°C and processed within the use by date. For raw chicken sold as raw dog food (RDF), one of each brand sold across five RDF stores in Bristol was purchased. A total of 15 packets were collected in September 2023, covering 11 brands of RDF.

Under sterile conditions, 200g of each meat/RDF sample was placed into a stomacher bag with 200 mL of EC broth (Oxoid ltd). Bags were sealed and incubated for 5 h at 37°C, 180 rpm. Each liquid culture was aliquoted into Eppendorf tubes containing sterile glycerol (to 25% v/v final) and stored at -70°C.

### 2.3. Microbiology

Twenty microlitres of each thawed liquid culture aliquot (above) was spread onto Tryptone Bile X-Glucuronide agar plates (Sigma) containing Nitrofurantoin (64 mg/L) based on EUCAST resistance breakpoints for human infections [24] and incubated overnight at 37°C. The media were quality control checked by plating known NFT-R and susceptible isolates. Putative NFT-R *E. coli* were re-streaked using the same medium to confirm resistance. Isolates were stored on cryobeads at -70°C. Resistance was confirmed by MIC testing using broth microdilution according to EUCAST methodology [24].

### 2.4. WGS and phylogenetic analysis

WGS was performed by MicrobesNG to achieve a minimum 30-fold coverage. Preparation of isolates was carried out following MicrobesNG strain submission procedures as described [25]. Genomes were sequenced on an Illumina NovaSeq 6000 (Illumina, San Diego, USA) using a 250 bp paired end protocol. Sequencing data were analysed using ResFinder 4.4.2 [26], Hound [27] and MLST 2.0 [28].

Sequence alignment and phylogenetic analysis was carried out using the computational facilities of the Advanced Computing Research Centre, University of Bristol (http://www.bristol.ac.uk/acrc/) and the Bioconda environment management system [29]. Alignment with reference sequences (**Table S1**) was performed using Snippy and Snippy-Core software (https://github.com/tseemann/snippy) and SNP distances were determined using SNP-dists (https://github.com/tseemann/snp-dists). Maximum likelihood phylogenetic trees were produced using RAxML and the GTRGAMMA model of rate of heterogeneity [30]. Phylogenetic trees were illustrated using Microreact [31]. Where only raw sequence reads were available on Enterobase [32], SPAdes [33] was used to assemble these sequences prior to inclusion into the phylogenetic analysis.

## 3. Results

### 3.1. NFT-R *E. coli* excreted by dogs being fed raw (uncooked) meat

We tested faecal samples from 297 healthy dogs living in the city of Bristol. Two dogs excreted NFT-R *E. coli* at detectable levels. The NFT-R *E. coli* were ST6805 and ST919 and both had loss-of-function mutations in *nfsA* and *nfsB* (**Table 1**).

**Table 1.** NFT-R *E. coli* isolates excreted by dogs, and from raw meat in Bristol, UK.

| Isolate | ST | Phylo-group | Effect of non-synonymous mutations relative to *E. coli* K12 |  |  |
| --- | --- | --- | --- | --- | --- |
|  |  |  | NfsA | NfsB | Other resistance genes |
| RCM1 | ST8874 | B2 | I117T, K141E, G187D, 212Stop | 46Stop | None |
| RCM2 | ST665 | A | 138FS | M75I, V93A, 101Stop | *aadA1*, *aph(3'')-Ib*, *aph(6)-Id*,<br>*sul1*, *sul2*, *bla*<sub>TEM-1B</sub>, *tet(A)* |
| RDF15 | ST8874 | B2 | I117T, K141E, G187D, 212Stop | 46Stop | None |
| DOG1030 | ST919 | B2 | I117T, K141E, G187D, 212Stop | 46Stop | None |
| DOG1110 | ST6805 | A | Insertional inactivation after codon 225 | M75I, 79Stop | *aadA5*, *dfrA17*, *sul2*, *tet(A)* |

Isolates were from raw chicken meat sold for human consumption (RCM), chicken raw dog food (RDF), and canine faeces (DOG), sequence type (ST) and missense and nonsense mutations, deletions and insertions affecting NfsA and NfsB sequences are noted. Green represents those variants predicted to be functionally wild-type. Red represents those conferring loss of function [2].

We collected owner-completed survey data about the lives of these 297 Bristol dogs [20,21]. Since only two dogs excreted NFT-R *E. coli* it is not possible to be confident about any statistical assessment of associations between excretion and any lifestyle practice. Nonetheless, we noted (**Table S2**) that both dogs were fed raw meat at the time of faecal sampling, which was a practice observed for only 31/297 dogs (Fisher Exact p=0.011).

### 3.2. NFT-R *E. coli* in uncooked chicken meat and chicken-based commercial RDF

In the UK, raw meat fed to dogs is sourced as commercial RDF or meat sold primarily for human consumption (once cooked) [34]. Accordingly, to investigate whether raw meat potentially being fed to dogs in the city of Bristol was contaminated with NFT-R *E. coli*, we tested 73 samples of meat from across the city. These were 15 packets each of beef, chicken and lamb, and 13 of pork purchased from large-chain grocery stores, and 15 packets of chicken-based commercial RDF of various brands.

Two out of 58 (3%) samples of meat from large-chain grocery stores tested positive for NFT-R *E. coli*; both were chicken meat. WGS showed that these two isolates had loss of function mutations in *nfsA* and *nfsB* and were ST665 and ST8874 (**Table 1**). Out of fifteen packets of chicken-based commercial RDF, one (7%) was positive for NFT-R *E. coli*. WGS showed this to be ST8874, the same ST found in one packet of chicken meat (**Table 1**).

### 3.3. Poultry-associated NFT-R and NFT pre-resistant *E. coli* clones found across multiple countries and One Health compartments, including human UTI

The NFT-R ST8874 and ST919 isolates found on meat or excreted by one dog in Bristol had the same mutations in *nfsA* and in *nfsB* and none of them carried acquired resistance genes (**Table 1**). These STs are single-locus variants: *adk*38 in ST8874 versus *adk*861 in ST919 but otherwise they share the same multi-locus ST alleles.

We found 71 ST919 single-locus variant genomes with available sequence data on Enterobase (search date, 2^nd^ May 2025) (**Table S3**). Of these, 16 carried *nfsA/B* mutations identical to the Bristol ST919/ST8874 isolates reported above, with the rest being wild-type (**Table S3**). One of the *nfsA/B* mutants was an ST8874 isolate from a chicken yolk sac infection in Czechia [35]. The other 15 *nfsA/B* mutants were ST919, of which one isolate is a published NFT-R *E. coli* from a human UTI from London, UK [6]. Of the remaining *nfsA/B* mutant ST919 genomes on Enterobase, all are from chicken caecal contents/faeces or chicken meat. They are from the UK, Canada, the USA and Japan (**Table S3**).

We performed a core genome alignment of the ST919/ST8874 *nfsA*/*B* mutant isolates and present a matrix of single nucleotide polymorphism (SNP) distances between pairs of isolates (**Table S4**) and a phylogenetic tree (**Fig. 1**). The two Bristol ST8874 meat isolates and the Czech yolk sac infection isolate were very closely related to each other (<60 SNPs across a core genome alignment), and to an ST919 UK poultry isolate on Enterobase (<100 SNPs) which was only 33 SNPs different from the NFT-R human UTI isolate from London, UK. The ST8874 clone appears to have branched off from this UK ST919 clone (**Fig. 1**). However, the ST919 isolate excreted by a dog in Bristol was more closely related to chicken isolates from the USA and Canada (49-73 SNPs) than to the ST919 UK chicken isolate (172 SNPs). The ST919 isolates from North America were variable in their relatedness but did noticeably cluster together (**Fig. 1**). The most diverse isolate was from Japan (**Table S4, Fig. 1**).

**Figure 1.**
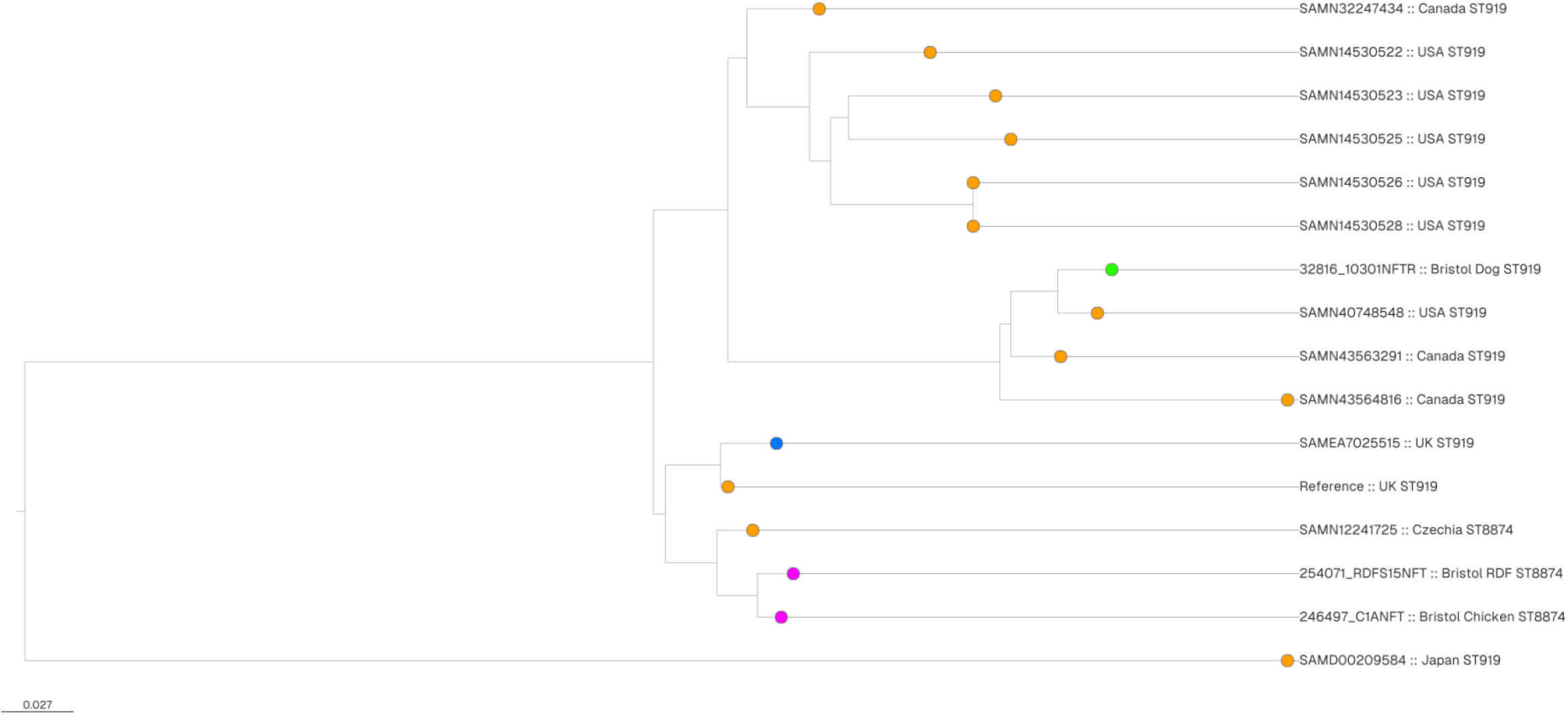
Maximum Likelihood phylogenetic tree based on a core genome alignment illustrating the evolutionary relationship between NFT-R ST919/ST8874 NfsA/NfsB mutant isolates. Core genomes from Enterobase isolates (from poultry sources, orange; from human sources, blue) and chicken meat/RDF (pink) or dogs (green) from Bristol were aligned using Snippy and Phylogenetic trees were constructed based on RaXML analysis. Isolate origins as reported in Enterobase and their STs are noted.

After establishing the existence of a poultry-associated global NFT-R ST919/ST8874 clone (**Fig. 1**), we next considered whether the ST665 and ST6805 NFT-R *E. coli* from meat and dogs in Bristol (**Table 1**) also cluster into widely dispersed clones. We found 95 ST665 sequences on Enterobase. Of 77 with a defined origin, 46 (60%) were from poultry meat or poultry farms and 12 (16%) from humans. Two of these human isolates and three isolates from chicken faeces, all from the UK had the same *nfsA* (138FS) and *nfsB* (101Stop) mutations as the NFT-R ST665 isolate from chicken meat in Bristol (**Table S5**).

Multiple human-derived ST665 isolates from across Europe and into the Middle East had loss-of-function mutations in *nfsA* but wild-type *nfsB* (**Table S5**) and so are considered nitrofurantoin pre-resistant [2]. In some cases this was the same 138FS *nfsA* mutation seen in the UK NFT-R isolates, with other common *nfsA* mutations being 46FS and 49FS. These same three mutations were observed among ST665 isolates from poultry and wild birds in multiple countries (**Table S5**).

A SNP distance analysis revealed that the six UK NFT-R *nfsA* (138FS), *nfsB* (101Stop) isolates are diverse. For example, the Bristol chicken meat isolate is 1773 SNPs away from the closest UK chicken faeces isolate and 2095 away from the closest UK human isolate. Furthermore, the chicken faeces *nfsA* (138FS), *nfsB* (101Stop) isolates were at least 1964 SNPs away from the human isolates (**Table S6**). This divergence was reflected in the resulting phylogenetic tree (**Fig. 2**). In other regards the isolates from Enterobase phylogenetically clustered based on their *nfsA* allele (**Fig. 2**). Indeed, it appears from this analysis that the NFT-R ST665 NFT-R *nfsA* (138FS), *nfsB* (101Stop) isolates have evolved from three divergent 138FS lineages (**Fig. 2**).

**Figure 2.**
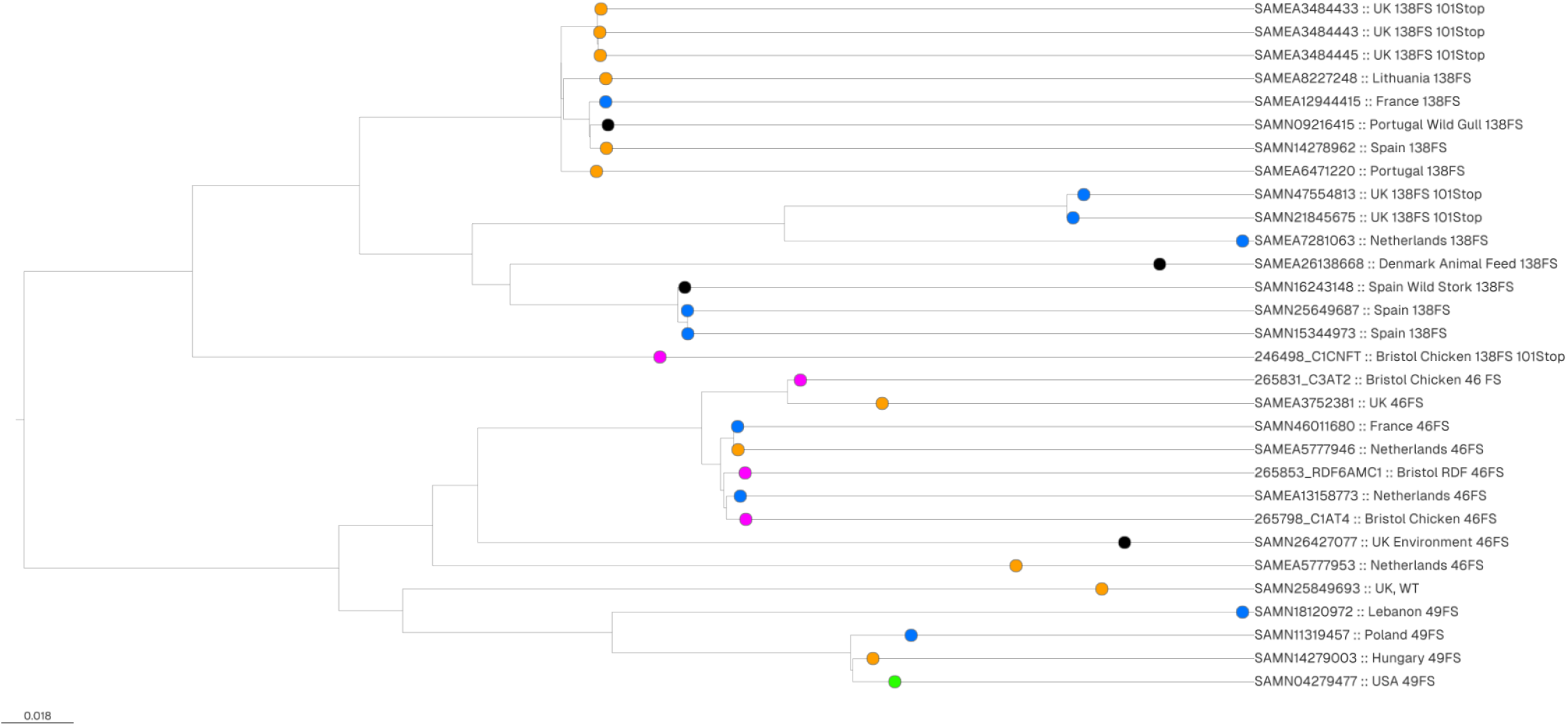
Maximum Likelihood phylogenetic tree based on a core genome alignment illustrating the evolutionary relationship between NFT-R and pre-resistant ST665 isolates. Core genomes from Enterobase isolates (from poultry sources, orange; from human sources, blue; from dogs, green; from other sources as noted, black) and chicken meat/RDF isolates from Bristol (pink) were aligned using Snippy and Phylogenetic trees were constructed based on RaXML analysis. Isolate origins as reported in Enterobase and the NfsA mutations (plus NfsB 101Stop where relevant) are noted.

For all three main ST665 nitrofurantoin pre-resistant lineages (46FS, 49FS or 138FS *nfsA* alleles but having wild-type *nfsB*) there are instances of likely sharing between poultry, wild birds and humans (**Table S6, Fig. 2**). For example, the 46FS human isolates from the Netherlands and France are 94 and 91 SNPs, respectively, different from a chicken isolate from the Netherlands. A Spanish human 138FS isolate was 42 SNPs different from an isolate from a wild Stork, also from Spain. Finally, a Polish human 49FS isolate was 207 SNPs different from a Hungarian chicken isolate.

Among four ST6805 isolates on Enterobase, one Danish poultry isolate (SAMEA112328101) shared the same *nfsA* mutation as a Bristol dog isolate (**Table 1**), but it had a wild type *nfsB*, hence is also considered pre-resistant. These two isolates are 2281 SNPs apart, however, so there is no evidence of a circulating clone.

Based on the above findings, we suspected that there might be additional nitrofurantoin pre-resistant clones present on chicken meat being sold in Bristol. To test this, we sequenced 92 nitrofurantoin susceptible *E. coli* from RDF and 76 from uncooked chicken meat, cultured from the same samples as the NFT-R isolates (above), but which did not grow on nitrofurantoin agar at a concentration required to define them as resistant. We found 17 isolates from 12/15 samples of chicken meat and 10 isolates from 4/15 samples of chicken-based RDF that had obvious loss of function in *nfsA* but wild-type *nfsB* and so are defined as pre-resistant. These isolates, accounting for 16.1% of all isolates from chicken meat or RDF sequenced, spanned 11 STs (**Table S7**). They included ST665 isolates from two chicken and one RDF samples with identical *nfsA* alleles (46FS). These had pairwise SNP distances ranging from 110 to 355, and are part of the widely disseminated 46FS pre-resistant clone, defined above, with one chicken and one RDF isolate being 84 and 96 SNPs different, respectively, from a human isolate from the Netherlands (**Table S6; Fig. 2**).

Pre-resistant ST69 isolates with identical *nfsA* mutations were identified from six chicken meat and one RDF samples collected in Bristol with SNP distances of 10 to 41 between them (**Table S7**). The other STs found to include *nfsA* mutants were ST10, ST57, ST58, ST155, ST162 and ST2705. Furthermore, for ST7529 and ST752 *nfsA* mutants found on the chicken meat/RDF in Bristol, we found overlap with human isolates. For ST7529, one human clinical Enterobase isolate from the UK was found to be 103 SNPs different from the two Bristol RDF isolates, which were themselves identical. All carried the same *nfsA* loss of function mutation (**Table S7**).

Of ST752 isolates on Enterobase (n=503), the majority were from poultry (33.0%) or human (28.2%) sources. Of these, 91.9% of human isolates and 84.9% of poultry isolates carried the same *nfsA* mutation as also seen in the Bristol chicken meat isolates suggesting they are nitrofurantoin pre-resistant. We selected ST752 *nfsA* mutant Enterobase isolates from poultry and humans (n=10 for each) for comparison with the six isolates from chicken meat in Bristol (**Table S7**). Two human isolates, one from the UK and the other from the Netherlands were 43 and 61 SNPs different, respectively, from one of the Bristol chicken meat isolates (**Table S7; Table S8**). Furthermore, there were four incidences where ST752 isolates from humans were <100 SNPs from Enterobase poultry isolates, with the closest pair being 48 SNPs apart (**Table S7; Table S8**).

## 4. Discussion

Because nitrofurantoin is not licenced for use in dogs in the UK, we were surprised to find NFT-R *E. coli* in faeces excreted by two dogs out of a cohort of 297 based on testing one faecal sample per dog with a limit of detection of approximately 1000 viable bacteria per gramme of faeces. The NFT-R *E. coli* from one dog is clonally related to those found in poultry meat and poultry farm environment samples in the UK, Canada, Czechia and Japan, including two single-locus variant STs (**Fig. 1**). Given that we also found NFT-R *E. coli* from this same clone in uncooked chicken meat, and chicken-based RDF purchased in the same city, a possible explanation for these dogs excreting poultry-associated NFT-R *E. coli* is the fact – reported by their owners – that they were being raw fed at the time of sampling. Whilst we have shown compelling evidence that feeding raw meat to dogs is associated with them excreting fluoroquinolone resistant *E. coli* [21], the evidence for NFT-R *E. coli* presented here is much weaker. This may be related to the relative degrees of contamination of meat with *E. coli* resistant to these different agents. For example, we identified here 10% (3/30) of chicken meat or chicken-based RDF samples were positive for NFT-R *E. coli*, but we have also reported that 47% (14/30) of these same samples were positive for fluoroquinolone-resistant *E. coli* [23].

Several studies have reported sharing of resistant *E. coli* between dogs and their owners [36-39], so it is possible that dogs excreting NFT-R *E. coli* may pass these on to their owners. This is in addition to any associated risks of accidental ingestion when handling and preparing RDF or other uncooked poultry, which we have shown here can be contaminated with numerous NFT-R or pre-resistant *E. coli* clones, particularly ST665 and ST752, which are also found in clinical samples from humans. Accordingly, it would seem prudent for those at risk of UTI, and particularly those who are taking nitrofurantoin, to be cautious when handling uncooked meat or interacting with pets fed uncooked meat, and practice enhanced hygiene regimes. This is also important for people taking other antibiotics, and particularly ciprofloxacin, but since the number of nitrofurantoin courses dispensed in the community in some countries is very high, and since resistance rates among urinary *E. coli* from humans are very low, the additional zoonotic burden exposed here is potentially important.

Whilst nitrofuran antibacterial use in poultry production was banned in many countries decades ago due to toxicity concerns if residues are found in meat [11], it is notable that SNP distances between isolates from the various NFT-R and pre-resistant clones reported here are large, possibly because they emerged many years ago.

It remains unclear how prevalent NFT-R or pre-resistant *E. coli* are in meat sold for human consumption throughout the UK and around the world, and whether other poultry-associated NFT-R or pre-resistant clones exist, but the novel findings of this study should stimulate interest in answering that question through widespread surveillance in multiple One Health compartments.

## Data Availability

Novel WGS data are deposited with ENA under accessions PRJEB74489 (Isolates from meat) and PRJEB74488 (isolates from canine faeces).

## Funding

This work was funded by grant NE/N01961X/1 to M.B.A. from the Antimicrobial Resistance Cross Council Initiative supported by the seven UK research councils. It was also funded by grant BB/X012670/1 from the Biotechnology and Biological Sciences Research Council. J.E.S. was supported by a scholarship from the Medical Research Foundation National PhD Training Programme in Antimicrobial Resistance Research (MRF-145-0004-TPG-AVISO).

## Author Contributions

Conceived the Study: J.E.S., M.B.A.

Collection of Data: J.E.S, B.A.

Cleaning and Analysis of Data: J.E.S. O.M.

Initial Drafting of Manuscript: J.E.S., M.B.A.

Corrected and Approved Manuscript: All authors

## Declaration of Competing Interest

M.B.A. is married to the owner of a veterinary practice that sells various mass-manufactured dog foods amounting to a value less than 5% of total turnover. Otherwise, none of the above named authors is declaring any conflict of interest in relation to the above titled submitted manuscript.

## Acknowledgements

Genome sequencing was provided by MicrobesNG (http://www.microbesng.uk). The authors thank Kate Rollings for assisting with data collection and all dog owners for agreeing to participate in this study.

## Notes

### Summary of Updates

Additional Enterobase genomes have become available and so the analysis has been expanded.

