## Supplementary Tables 1-8 for "Poultry-Associated Nitrofurantoin-Resistant and Pre-Resistant *Escherichia coli* Clones are Found in Multiple Countries and One-Health Compartments"

**Table S1 Reference genomes used for sequencing alignment**

| ST | ***E. coli* reference genome accession** |
|---|---|
| ST69 | SAMN40242201 |
| ST665 | SAMN25849693 |
| ST919 | SAMN11444779 |
| ST752 | SAMN32868256 |
| ST6805 | SAMEA112328101 |
| ST7529 | SAMN28864606 |

**Table S2. Practices common to both dogs excreting NFT-R *E. coli* from the city of Bristol as reported by owners in a survey compared with how common each practice is among the canine cohort (n=297) in this study**

| Practice | Positive for practice – NFT-R excreters | Positive for practice – Not NFT-R excreters | Fisher's exact (p=) |
|---|---|---|---|
| Often/very often exercising on roads/streets | 2/297 | 191/297 | 0.54 |
| Often/very often exercising in parks | 2/297 | 284/297 | 1 |
| Sometimes exercising on beaches | 2/297 | 195/297 | 0.5 |
| Sharing a home with a cat | 2/297 | 48/297 | 0.028 |
| Fed raw meat | 2/297 | 29/297 | 0.011 |

Practices positive for one NFT-R excreting dog only: walking in the countryside (not around livestock); swimming in lakes; swimming in ponds; swimming in rivers; received antibiotics in the last 6 months. Practices for neither NFT-R excreting dog: walking in the countryside around livestock; living with another animal except a cat,

#### Table S3. ST919 single-locus variant *E. coli* identified on Enterobase showing *nfsAB* mutation status and origins

| Sample Accession | Source | Location | ST | *nfsAB* mutation? |
|---|---|---|---|---|
| SAMN46723104 | | United States | 1865 | N |
| SAMN45102163 | | United States | 1865 | N |
| SAMN44490190 | | United States | 1865 | N |
| SAMN43564816 | Chicken Meat | Canada | 919 | Y |
| SAMN43563291 | Chicken Meat | Canada | 919 | Y |
| SAMN43303518 | Companion Animal | United States | 919 | N |
| SAMN41629489 | Human | United Kingdom | 919 | N |
| SAMN40748560 | Poultry | United States | 1333 | N |
| SAMN40748548 | Chicken Meat | United States | 919 | Y |
| SAMN40737978 | Companion Animal | United States | 919 | N |
| SAMN40372514 | Wild Animal | Italy | 1333 | N |
| SAMN36864136 | Companion Animal | United States | 1333 | N |
| SAMN32247434 | Chicken Caecal Contents | Canada | 919 | Y |
| SAMN29767770 | | United States | 1333 | N |
| SAMN29598476 | Companion Animal | United States | 5328 | N |
| SAMN29503068 | Companion Animal | Canada | 919 | N |
| SAMN29042213 | Poultry | United States | 919 | N |
| SAMN26305157 | Human | United States | 1333 | N |
| SAMN25142514 | Poultry | United States | 919 | N |
| SAMN20285927 | Human | United Kingdom | 1865 | N |
| SAMN19949776 | Wild Animal | United States | 919 | N |
| SAMN19069014 | Wild Animal | United States | 1333 | N |
| SAMN14593721 | Environment | United States | 919 | N |
| SAMN14530528 | Chicken Caecal Contents | United States | 919 | Y |
| SAMN14530526 | Chicken Caecal Contents | United States | 919 | Y |
| SAMN14530525 | Chicken Caecal Contents | United States | 919 | Y |
| SAMN14530524 | Chicken Caecal Contents | United States | 919 | Y |
| SAMN14530523 | Chicken Caecal Contents | United States | 919 | Y |
| SAMN14530522 | Chicken Caecal Contents | United States | 919 | Y |
| SAMN14113844 | Environment | United States | 919 | N |
| SAMN14113834 | Environment | United States | 919 | N |
| SAMN13721833 | Poultry | United States | 919 | N |
| SAMN13513936 | Environment | United States | 919 | N |
| SAMN13513935 | Environment | United States | 919 | N |
| SAMN13513926 | Environment | United States | 10536 | N |
| SAMN12359555 | Poultry | United States | 919 | N |
| SAMN12241725 | Chicken Yolk Sac Infection | Czechia | 8874 | Y |
| SAMN11444779 | Chicken Caecal Contents | United Kingdom | 919 | Y |
| SAMN10875599 | Human | Canada | 919 | N |
| SAMN10524481 | Human | United Kingdom | 1333 | N |
| SAMN10396941 | Environment | United States | 1865 | N |
| SAMN10221563 | Poultry | United States | 919 | N |
| SAMN08039647 | Environment | | 5328 | N |
| SAMN08039645 | Environment | | 5328 | N |
| SAMN08039644 | Environment | | 5328 | N |
| SAMN07678444 | Food | United States | 919 | N |
| SAMN07677607 | Food | United States | 919 | N |
| SAMN07677496 | Food | United States | 919 | N |
| SAMN05468056 | Livestock | United States | 919 | N |
| SAMN05464555 | Wild Animal | United States | 1333 | N |
| SAMN04992253 | Livestock | United States | 919 | N |
| SAMEA7025515 | Human UTI | United Kingdom | 919 | Y |
| SAMEA6163430 | Human | | 1333 | N |
| SAMEA4428396 | Human | Netherlands | 1333 | N |
| SAMEA115667394 | Human | Germany | 1333 | N |
| SAMEA115626452 | Wild Animal | Denmark | 919 | N |
| SAMEA115372530 | | United Kingdom | 919 | N |
| SAMEA114461330 | | | 11823 | N |
| SAMEA114461293 | | | 11823 | N |
| SAMEA114461292 | | | 11823 | N |
| SAMEA114461159 | | | 11823 | N |
| SAMEA104437735 | | Poland | 919 | N |
| SAMEA104437717 | | Poland | 919 | N |
| SAMEA104314597 | | | 919 | N |
| SAMEA104314596 | | | 919 | N |
| SAMEA104314595 | | | 919 | N |
| SAMEA104314589 | | | 1865 | N |
| SAMEA104027456 | | | 919 | N |
| SAMD00209584 | Chicken Caecal Contents | Japan | 919 | Y |
| SAMD00209572 | Chicken Caecal Contents | Japan | 919 | Y |
| SAMD00209571 | Chicken Caecal Contents | Japan | 919 | Y |

#### Table S4 SNP distance of ST919/ST8874 isolates carrying NfsA (212Stop) and NfsB (46Stop) loss-of-function mutations

| | | 246497_ C1ANFT | 254071_ RDFS15NFT | 32816_ 10301NFTR | Reference | SAMD 00209584 | SAMEA 7025515 | SAMN 12241725 | SAMN 14530522 | SAMN 14530523 | SAMN 14530525 | SAMN 14530526 | SAMN 32247434 | SAMN 40748548 | SAMN 43563291 | SAMN 43564816 |
|---|---|---|---|---|---|---|---|---|---|---|---|---|---|---|---|---|
| 246497_C1ANFT | Bristol. Chicken ST8874 | 0 | 31 | 193 | 91 | 514 | 112 | 49 | 175 | 197 | 204 | 192 | 143 | 186 | 168 | 371 |
| 254071_RDFS15NFT | Bristol. RDF ST8874 | 31 | 0 | 202 | 100 | 523 | 121 | 58 | 184 | 206 | 213 | 201 | 152 | 195 | 177 | 380 |
| 32816_10301NFTR | Bristol. Dog ST919 | 193 | 202 | 0 | 172 | 533 | 193 | 180 | 176 | 196 | 199 | 181 | 146 | 49 | 73 | 276 |
| SAMN11444779 (Reference) | UK. Chicken ST919 | 91 | 100 | 172 | 0 | 491 | 33 | 78 | 154 | 176 | 183 | 171 | 122 | 165 | 147 | 350 |
| SAMD00209584* | Japan. Chicken ST919 | 514 | 523 | 533 | 491 | 0 | 512 | 501 | 557 | 565 | 572 | 552 | 525 | 528 | 510 | 696 |
| SAMEA7025515 | UK. Human UTI ST919 | 112 | 121 | 193 | 33 | 512 | 0 | 99 | 175 | 197 | 204 | 192 | 143 | 186 | 168 | 371 |
| SAMN12241725 | Czechia. Chicken ST8874 | 49 | 58 | 180 | 78 | 501 | 99 | 0 | 162 | 184 | 191 | 179 | 130 | 173 | 155 | 358 |
| SAMN14530522 | USA. Chicken ST919 | 175 | 184 | 176 | 154 | 557 | 175 | 162 | 0 | 108 | 115 | 107 | 106 | 169 | 151 | 352 |
| SAMN14530523 | USA. Chicken ST919 | 197 | 206 | 196 | 176 | 565 | 197 | 184 | 108 | 0 | 125 | 119 | 128 | 189 | 171 | 366 |
| SAMN14530525 | USA. Chicken ST919 | 204 | 213 | 199 | 183 | 572 | 204 | 191 | 115 | 125 | 0 | 128 | 135 | 192 | 174 | 375 |
| SAMN14530526* | USA. Chicken ST919 | 192 | 201 | 181 | 171 | 552 | 192 | 179 | 107 | 119 | 128 | 0 | 123 | 174 | 156 | 353 |
| SAMN32247434 | Canada. Chicken ST919 | 143 | 152 | 146 | 122 | 525 | 143 | 130 | 106 | 128 | 135 | 123 | 0 | 139 | 121 | 324 |
| SAMN40748548 | USA. Chicken ST919 | 186 | 195 | 49 | 165 | 528 | 186 | 173 | 169 | 189 | 192 | 174 | 139 | 0 | 66 | 269 |
| SAMN43563291 | Canada. Chicken ST919 | 168 | 177 | 73 | 147 | 510 | 168 | 155 | 151 | 171 | 174 | 156 | 121 | 66 | 0 | 247 |
| SAMN43564816 | Canada. Chicken ST919 | 371 | 380 | 276 | 350 | 696 | 371 | 358 | 352 | 366 | 375 | 353 | 324 | 269 | 247 | 0 |

*These isolates are each representative of three identical isolates (Zero SNPs different)

**Table S5. Nitrofurantoin pre-resistant ST665 *E. coli* identified on Enterobase with obvious loss of function mutations in *nfsA***

| Accession | Country | Origin | NfsA variant | NfsB variant |
|---|---|---|---|---|
| SAMEA12944415 | France | HUMAN | 138FS | - |
| SAMEA26138668 | Denmark | Animal Feed | 138FS | - |
| SAMEA3484433 | UK | Chicken Faeces | 138FS | 101Stop |
| SAMEA3484443 | UK | Chicken Faeces | 138FS | 101Stop |
| SAMEA3484445 | UK | Chicken Faeces | 138FS | 101Stop |
| SAMEA6471220 | Portugal | POULTRY | 138FS | - |
| SAMEA7281063 | Netherlands | HUMAN | 138FS | - |
| SAMEA8227248 | Lithuania | POULTRY | 138FS | - |
| SAMN09216415 | Portugal | Wild Gull Faeces | 138FS | - |
| SAMN14278962 | Spain | POULTRY | 138FS | - |
| SAMN15344973 | Spain | HUMAN | 138FS | - |
| SAMN16243148 | Spain | White Stork | 138FS | - |
| SAMN21845675 | UK | HUMAN | 138FS | 101Stop |
| SAMN25649687 | Spain | HUMAN | 138FS | - |
| SAMN47554813 | UK | HUMAN | 138FS | 101Stop |
| SAMEA13158773 | Netherlands | HUMAN | 46FS | - |
| SAMEA3752381 | UK | Chicken Faeces | 46FS | - |
| SAMEA5777946 | Netherlands | Chicken Meat | 46FS | - |
| SAMEA5777953 | Netherlands | Chicken Meat | 46FS | - |
| SAMN26427077 | UK | Environment | 46FS | - |
| SAMN46011680 | France | HUMAN | 46FS | - |
| SAMN09580040 | Australia | Duck Faeces | 45Stop | - |
| SAMN04279477 | USA | Dog Faeces | 49FS | - |
| SAMN11319457 | Poland | HUMAN | 49FS | - |
| SAMN14279003 | Hungary | POULTRY | 49FS | - |
| SAMN18120972 | Lebanon | HUMAN | 49FS | - |
| SAMN21214633 | Italy | POULTRY | 68FS | - |
| SAMN48012225 | USA | Poultry | 100Stop | - |
| SAMN07678428 | USA | Unspecified Food | 175FS | - |
| SAMN26427274 | UK | Environment | - | 35FS |

### Table S6. SNP distances between ST665 NFT-R/pre-resistant isolates with NfsA and NfsB (as stated) mutations

| | | 246498_ C1CNFT | 265798_ C1AT4 | 265831_ C3AT2 | 265853_ RDF6AMC1 | SAMEA 5777946 | SAMEA 5777953 | SAMEA 13158773 | Reference | SAMEA 12944415 | SAMEA 26138668 | SAMEA 3484433 | SAMEA 3484443 | SAMEA 3484445 | SAMEA 3752381 | SAMEA 6471220 | SAMEA 7281063 | SAMEA 8227248 | SAMN 04279477 | SAMN 09216415 | SAMN 11319457 | SAMN 14278962 | SAMN 14279003 | SAMN 15344973 | SAMN 16243148 | SAMN 18120972 | SAMN 21845675 | SAMN 25649687 | SAMN 26427077 | SAMN 46011680 | SAMN 47554813 |
|---|---|---|---|---|---|---|---|---|---|---|---|---|---|---|---|---|---|---|---|---|---|---|---|---|---|---|---|---|---|---|---|
| 246498_ C1CNFT | Bristol Chicken 138FS 101Stop | 0 | 2836 | 2969 | 2836 | 2818 | 2693 | 2824 | 3071 | 1966 | 2912 | 1776 | 1773 | 1774 | 2873 | 1943 | 2645 | 1966 | 2932 | 1972 | 2930 | 1968 | 2915 | 1921 | 1914 | 3498 | 2095 | 1920 | 3030 | 2817 | 2120 |
| 265798_ C1AT4 | Bristol Chicken 46FS | 2836 | 0 | 355 | 110 | 108 | 1861 | 84 | 2365 | 1856 | 3743 | 1844 | 1841 | 1842 | 553 | 1817 | 3095 | 1856 | 1849 | 1862 | 1852 | 1858 | 1830 | 2864 | 2857 | 2545 | 3129 | 2863 | 2001 | 105 | 3154 |
| 265831_ C3AT2 | Bristol Chicken 46FS | 2969 | 355 | 0 | 353 | 335 | 2088 | 341 | 2484 | 1991 | 3602 | 1975 | 1972 | 1973 | 268 | 1952 | 3230 | 1975 | 1973 | 1997 | 1973 | 1993 | 1951 | 2999 | 2992 | 2658 | 3260 | 2998 | 1768 | 334 | 3285 |
| 265853_ RDF6AMC1 | Bristol RDF 46FS | 2836 | 110 | 353 | 0 | 106 | 1861 | 96 | 2365 | 1854 | 3741 | 1842 | 1839 | 1840 | 551 | 1815 | 3093 | 1854 | 1849 | 1860 | 1852 | 1856 | 1830 | 2862 | 2855 | 2545 | 3127 | 2861 | 1999 | 105 | 3152 |
| SAMEA 5777946 | Netherlands Chicken 46FS | 2818 | 108 | 335 | 106 | 0 | 1841 | 94 | 2347 | 1836 | 3723 | 1824 | 1821 | 1822 | 533 | 1797 | 3075 | 1836 | 1831 | 1842 | 1834 | 1838 | 1812 | 2844 | 2837 | 2527 | 3109 | 2843 | 1981 | 21 | 3134 |
| SAMEA 5777953 | Netherlands Chicken 46FS | 2693 | 1861 | 2088 | 1861 | 1841 | 0 | 1849 | 2843 | 3203 | 4145 | 3191 | 3188 | 3189 | 2286 | 3164 | 3631 | 3203 | 2311 | 3209 | 2314 | 3205 | 2292 | 3133 | 3126 | 2829 | 3411 | 3132 | 2523 | 1840 | 3436 |
| SAMEA 13158773 | Netherlands Human 46FS | 2824 | 84 | 341 | 96 | 94 | 1849 | 0 | 2353 | 1842 | 3729 | 1830 | 1827 | 1828 | 539 | 1803 | 3081 | 1842 | 1837 | 1848 | 1840 | 1844 | 1818 | 2850 | 2843 | 2533 | 3115 | 2849 | 1987 | 91 | 3140 |
| Reference | | 3071 | 2365 | 2484 | 2365 | 2347 | 2843 | 2353 | 0 | 3204 | 4154 | 3192 | 3189 | 3190 | 2682 | 3165 | 3535 | 3132 | 2463 | 3210 | 2477 | 3206 | 2412 | 3234 | 3227 | 2909 | 3472 | 3233 | 2813 | 2346 | 3497 |
| SAMEA 12944415 | France Human 138FS | 1966 | 1856 | 1991 | 1854 | 1836 | 3203 | 1842 | 3204 | 0 | 2150 | 212 | 209 | 210 | 1895 | 199 | 2232 | 212 | 2794 | 88 | 2792 | 84 | 2777 | 1314 | 1307 | 3420 | 2153 | 1313 | 2782 | 1835 | 2178 |
| SAMEA 26138668 | Denmark Animal Feed 138FS | 2912 | 3743 | 3602 | 3741 | 3723 | 4145 | 3729 | 4154 | 2150 | 0 | 2139 | 2136 | 2137 | 3506 | 2110 | 3156 | 2135 | 3907 | 2156 | 3902 | 2152 | 3887 | 1810 | 1803 | 4446 | 2603 | 1809 | 3504 | 3722 | 2628 |
| SAMEA 3484433 | UK Chicken 138FS 101Stop | 1776 | 1844 | 1975 | 1842 | 1824 | 3191 | 1830 | 3192 | 212 | 2139 | 0 | 15 | 16 | 1879 | 189 | 2220 | 212 | 2786 | 218 | 2784 | 214 | 2769 | 1302 | 1295 | 3412 | 1967 | 1301 | 2766 | 1823 | 1992 |
| SAMEA 3484443 | UK Chicken 138FS 101Stop | 1773 | 1841 | 1972 | 1839 | 1821 | 3188 | 1827 | 3189 | 209 | 2136 | 15 | 0 | 9 | 1876 | 186 | 2217 | 209 | 2783 | 215 | 2781 | 211 | 2766 | 1299 | 1292 | 3409 | 1964 | 1298 | 2763 | 1820 | 1989 |
| SAMEA 3484445 | UK Chicken 138FS 101Stop | 1774 | 1842 | 1973 | 1840 | 1822 | 3189 | 1828 | 3190 | 210 | 2137 | 16 | 9 | 0 | 1877 | 187 | 2218 | 210 | 2784 | 216 | 2782 | 212 | 2767 | 1300 | 1293 | 3410 | 1965 | 1299 | 2764 | 1821 | 1990 |
| SAMEA 3752381 | UK Chicken 46FS | 2873 | 553 | 268 | 551 | 533 | 2286 | 539 | 2682 | 1895 | 3506 | 1879 | 1876 | 1877 | 0 | 1856 | 3134 | 1879 | 2171 | 1901 | 2171 | 1897 | 2149 | 2903 | 2896 | 2856 | 3164 | 2902 | 1966 | 532 | 3189 |
| SAMEA 6471220 | Portugal Chicken 138FS | 1943 | 1817 | 1952 | 1815 | 1797 | 3164 | 1803 | 3165 | 199 | 2110 | 189 | 186 | 187 | 1856 | 0 | 2193 | 201 | 2757 | 205 | 2755 | 201 | 2740 | 1161 | 1154 | 3383 | 2130 | 1160 | 2743 | 1796 | 2155 |
| SAMEA 7281063 | Netherlands Human 138FS | 2645 | 3095 | 3230 | 3093 | 3075 | 3631 | 3081 | 3535 | 2232 | 3156 | 2220 | 2217 | 2218 | 3134 | 2193 | 0 | 2232 | 3567 | 2238 | 3565 | 2234 | 3550 | 2152 | 2145 | 4132 | 1654 | 2151 | 3364 | 3074 | 1679 |
| SAMEA 8227248 | Lithuania Chicken 138FS | 1966 | 1856 | 1975 | 1854 | 1836 | 3203 | 1842 | 3132 | 212 | 2135 | 212 | 209 | 210 | 1879 | 201 | 2232 | 0 | 2766 | 218 | 2724 | 214 | 2715 | 1316 | 1309 | 3361 | 2153 | 1315 | 2766 | 1835 | 2178 |
| SAMN 04279477 | USA Dog 49FS | 2932 | 1849 | 1973 | 1849 | 1831 | 2311 | 1837 | 2463 | 2794 | 3907 | 2786 | 2783 | 2784 | 2171 | 2757 | 3567 | 2766 | 0 | 2800 | 262 | 2796 | 155 | 2995 | 2988 | 1976 | 3329 | 2994 | 2686 | 1830 | 3350 |
| SAMN 09216415 | Portugal Gull 138FS | 1972 | 1862 | 1997 | 1860 | 1842 | 3209 | 1848 | 3210 | 88 | 2156 | 218 | 215 | 216 | 1901 | 205 | 2238 | 218 | 2800 | 0 | 2798 | 86 | 2783 | 1320 | 1313 | 3426 | 2159 | 1319 | 2788 | 1841 | 2184 |
| SAMN 11319457 | Poland Human 49FS | 2930 | 1852 | 1973 | 1852 | 1834 | 2314 | 1840 | 2477 | 2792 | 3902 | 2784 | 2781 | 2782 | 2171 | 2755 | 3565 | 2724 | 262 | 2798 | 0 | 2794 | 207 | 2993 | 2986 | 2021 | 3327 | 2992 | 2686 | 1833 | 3352 |
| SAMN 14278962 | Spain Chicken 138FS | 1968 | 1858 | 1993 | 1856 | 1838 | 3205 | 1844 | 3206 | 84 | 2152 | 214 | 211 | 212 | 1897 | 201 | 2234 | 214 | 2796 | 86 | 2794 | 0 | 2779 | 1316 | 1309 | 3422 | 2155 | 1315 | 2784 | 1837 | 2180 |
| SAMN 14279003 | Hungary Chicken 49FS | 2915 | 1830 | 1951 | 1830 | 1812 | 2292 | 1818 | 2412 | 2777 | 3887 | 2769 | 2766 | 2767 | 2149 | 2740 | 3550 | 2715 | 155 | 2783 | 207 | 2779 | 0 | 2978 | 2971 | 1933 | 3312 | 2977 | 2664 | 1811 | 3337 |
| SAMN 15344973 | Spain Human 138FS | 1921 | 2864 | 2999 | 2862 | 2844 | 3133 | 2850 | 3234 | 1314 | 1810 | 1302 | 1299 | 1300 | 2903 | 1161 | 2152 | 1316 | 2995 | 1320 | 2993 | 1316 | 2978 | 0 | 43 | 3452 | 1757 | 1 | 2842 | 2843 | 1782 |
| SAMN 16243148 | Spain Stork 138FS | 1914 | 2857 | 2992 | 2855 | 2837 | 3126 | 2843 | 3227 | 1307 | 1803 | 1295 | 1292 | 1293 | 2896 | 1154 | 2145 | 1309 | 2988 | 1313 | 2986 | 1309 | 2971 | 43 | 0 | 3445 | 1750 | 42 | 2835 | 2836 | 1775 |
| SAMN 18120972 | Lebanon Human 49FS | 3498 | 2545 | 2658 | 2545 | 2527 | 2829 | 2533 | 2909 | 3420 | 4446 | 3412 | 3409 | 3410 | 2856 | 3383 | 4132 | 3361 | 1976 | 3426 | 2021 | 3422 | 1933 | 3452 | 3445 | 0 | 3872 | 3451 | 3274 | 2526 | 3897 |
| SAMN 21845675 | UK Human 138FS 101Stop | 2095 | 3129 | 3260 | 3127 | 3109 | 3411 | 3115 | 3472 | 2153 | 2603 | 1967 | 1964 | 1965 | 3164 | 2130 | 1654 | 2153 | 3329 | 2159 | 3327 | 2155 | 3312 | 1757 | 1750 | 3872 | 0 | 1756 | 3297 | 3108 | 59 |
| SAMN 25649687 | Spain Human 138FS | 1920 | 2863 | 2998 | 2861 | 2843 | 3132 | 2849 | 3233 | 1313 | 1809 | 1301 | 1298 | 1299 | 2902 | 1160 | 2151 | 1315 | 2994 | 1319 | 2992 | 1315 | 2977 | 1 | 42 | 3451 | 1756 | 0 | 2841 | 2842 | 1781 |
| SAMN 26427077 | UK Environment 46FS | 3030 | 2001 | 1768 | 1999 | 1981 | 2523 | 1987 | 2813 | 2782 | 3504 | 2766 | 2763 | 2764 | 1966 | 2743 | 3364 | 2766 | 2686 | 2788 | 2686 | 2784 | 2664 | 2842 | 2835 | 3274 | 3297 | 2841 | 0 | 1980 | 3322 |
| SAMN 46011680 | France Human 46FS | 2817 | 105 | 334 | 105 | 21 | 1840 | 91 | 2346 | 1835 | 3722 | 1823 | 1820 | 1821 | 532 | 1796 | 3074 | 1835 | 1830 | 1841 | 1833 | 1837 | 1811 | 2843 | 2836 | 2526 | 3108 | 2842 | 1980 | 0 | 3133 |
| SAMN 47554813 | UK Human 138FS 101Stop | 2120 | 3154 | 3285 | 3152 | 3134 | 3436 | 3140 | 3497 | 2178 | 2628 | 1992 | 1989 | 1990 | 3189 | 2155 | 1679 | 2178 | 3350 | 2184 | 3352 | 2180 | 3337 | 1782 | 1775 | 3897 | 59 | 1781 | 3322 | 3133 | 0 |

**Table S7. Nitrofurantoin pre-resistant isolates from chicken meat and RDF in Bristol, United Kingdom, and relationships with human clinical isolates from Enterobase.**

| ST | Sample Type | NfsA Variant (all loss of function) | NfsB Variant | Human isolates with identical *nfsA* mutation (<150 SNPs) |
|---|---|---|---|---|
| 10 | Chicken Meat | 226 Ins IS*1* | WT | None |
| 57 | Chicken Meat | M1I (ATG-ATA) | V93A | None |
| 58 | RDF | 141 Ins IS*1* | M75I V93A | None |
| 69 | Chicken Meat (6 samples) | M1I | V93A | None |
| 69 | RDF | M1I | V93A | None |
| 155 | RDF | Del <150nt – 26 | M75I V93A P209L | None |
| 162 | RDF | Del 109-110 | M75I V93A A169T | None |
| 665 | Chicken Meat (2 samples) | 46FS | M75I V93A | SAMEA13158773 (Netherlands), 84 SNPs SAMN46011680 (France), 105 SNPs |
| 665 | RDF | 46FS | M75I V93A | SAMEA13158773 (Netherlands), 96 SNPs SAMN46011680 (France), 105 SNPs |
| 752 | Chicken Meat (5 samples) | M1I | WT | SAMN32868256 (UK), 43 SNPs SAMN25850013 (Netherlands), 61 SNPs SAMN14734242 (South Korean), 101 SNPs SAMN22183865 (UK), 135 SNPs |
| 2509 | RDF | 44Stop | M75I V93A | None |
| 2705 | Chicken Meat | 100Stop | M78I H80L V93A | None |
| 7529 | RDF (2 samples) | 88Stop | M75I V93A | SAMN092904 (UK), 103 SNPs |

### Table S8 SNP distance of ST752 isolates

| SampleID | Source | Country | Date | 236996_C2 Cspec | 245209_C5 Cspec | 245223_C4 Camox | 265837_C3 CT3 | 265839_C4 AT1 | 265840_C4 AT2 | SAMN3223 5060 | SAMN4044 2795 | SAMN4138 1106 | SAMN1084 0056 | SAMEA757 8064 | SAMEA757 8177 | SAMEA822 7255 | SAMEA822 7341 | SAMEA129 43667 | SAMN1144 4790 | SAMN3286 8256* | SAMN2218 3865 | SAMN3090 0011 | SAMN4023 1774 | SAMN4049 7913 | SAMN4144 9061 | SAMN1473 4242 | SAMN3041 6431 | SAMEA363 8314 | SAMN2585 0018 | SAMN2585 0013 |
|---|---|---|---|---|---|---|---|---|---|---|---|---|---|---|---|---|---|---|---|---|---|---|---|---|---|---|---|---|---|---|
| SAMN32868256* | Human | United Kingdom | 2023 | 2707 | 2785 | 43 | 2703 | 2709 | 1840 | 117 | 116 | 2004 | 2017 | 1147 | 48 | 2242 | 2343 | 109 | 2311 | 0 | 144 | 2015 | 3528 | 580 | 2602 | 110 | 2863 | 2588 | 3126 | 70 |
| SAMN22183865 | Human | United Kingdom | 2021 | 2733 | 2811 | 135 | 2729 | 2735 | 1864 | 113 | 112 | 2002 | 2041 | 1171 | 122 | 2266 | 2365 | 105 | 2335 | 144 | 0 | 2039 | 3554 | 604 | 2630 | 144 | 2887 | 2610 | 3154 | 162 |
| SAMN30900011 | Human | United Kingdom | 2022 | 2424 | 2508 | 2006 | 2426 | 2430 | 674 | 2012 | 2013 | 2535 | 52 | 2360 | 1993 | 2074 | 2403 | 2006 | 2281 | 2015 | 2039 | 0 | 3759 | 2190 | 3375 | 2017 | 2999 | 2811 | 3193 | 2033 |
| SAMN40231774 | Human | United Kingdom | 2023 | 3360 | 3466 | 3519 | 3384 | 3390 | 3379 | 3529 | 3526 | 3398 | 3761 | 4165 | 3506 | 3242 | 3563 | 3521 | 3798 | 3528 | 3554 | 3759 | 0 | 3689 | 4886 | 3532 | 3918 | 1351 | 3929 | 3546 |
| SAMN40497913 | Human | United Kingdom | 2024 | 2858 | 2936 | 571 | 2854 | 2860 | 2189 | 577 | 576 | 2111 | 2192 | 1273 | 558 | 2226 | 2314 | 569 | 2323 | 580 | 604 | 2190 | 3689 | 0 | 2706 | 580 | 2913 | 2755 | 3208 | 598 |
| SAMN41449061 | Human | United Kingdom | 2023 | 3870 | 4006 | 2593 | 3924 | 3930 | 3442 | 2603 | 2602 | 3371 | 3377 | 3177 | 2580 | 3351 | 3494 | 2595 | 2733 | 2602 | 2630 | 3375 | 4886 | 2706 | 0 | 2603 | 4049 | 3979 | 4327 | 2620 |
| SAMN14734242 | Human | South Korea | 2018 | 2709 | 2787 | 101 | 2705 | 2711 | 1842 | 117 | 116 | 2004 | 2019 | 1147 | 88 | 2242 | 2343 | 109 | 2311 | 110 | 144 | 2017 | 3532 | 580 | 2603 | 0 | 2863 | 2588 | 3128 | 126 |
| SAMN30416431 | Human | Ecuador | 2021 | 3131 | 3199 | 2854 | 3117 | 3123 | 2962 | 2860 | 2859 | 2647 | 3001 | 2388 | 2841 | 3130 | 2583 | 2852 | 2834 | 2863 | 2887 | 2999 | 3918 | 2913 | 4049 | 2863 | 0 | 3052 | 3722 | 2881 |
| SAMEA3638314 | Human | Germany | 2006 | 2426 | 2523 | 2579 | 2441 | 2447 | 2435 | 2585 | 2584 | 2572 | 2813 | 3221 | 2566 | 2295 | 2680 | 2577 | 3023 | 2588 | 2610 | 2811 | 1351 | 2755 | 3979 | 2588 | 3052 | 0 | 3052 | 2606 |
| SAMN25850018 | Human | Netherlands | 2021 | 3377 | 3415 | 3117 | 3333 | 3339 | 3091 | 3127 | 3126 | 3338 | 3195 | 3773 | 3104 | 2938 | 3221 | 3119 | 3116 | 3126 | 3154 | 3193 | 3929 | 3208 | 4327 | 3128 | 3722 | 3052 | 0 | 3144 |
| SAMN25850013 | Human | Netherlands | 2021 | 2725 | 2803 | 61 | 2721 | 2727 | 1858 | 135 | 132 | 2022 | 2035 | 1165 | 66 | 2260 | 2361 | 127 | 2329 | 70 | 162 | 2033 | 3546 | 598 | 2620 | 126 | 2881 | 2606 | 3144 | 0 |
| SAMN32235060 | Poultry | United States | 2022 | 2704 | 2782 | 108 | 2700 | 2706 | 1837 | 0 | 47 | 1957 | 2014 | 1144 | 95 | 2239 | 2340 | 64 | 2306 | 117 | 113 | 2012 | 3529 | 577 | 2603 | 117 | 2860 | 2585 | 3127 | 135 |
| SAMN40442795 | Poultry | Mexico | 2017 | 2705 | 2783 | 107 | 2701 | 2707 | 1838 | 47 | 0 | 1956 | 2015 | 1143 | 94 | 2238 | 2339 | 63 | 2307 | 116 | 112 | 2013 | 3526 | 576 | 2602 | 116 | 2859 | 2584 | 3126 | 132 |
| SAMN41381106 | Poultry | United States | 2024 | 2843 | 2908 | 1995 | 2826 | 2832 | 2432 | 1957 | 1956 | 0 | 2537 | 2620 | 1982 | 2348 | 2126 | 1953 | 2353 | 2004 | 2002 | 2535 | 3398 | 2111 | 3371 | 2004 | 2647 | 2572 | 3338 | 2022 |
| SAMN10840056 | Poultry | United Kingdom | 2015 | 2426 | 2510 | 2008 | 2428 | 2432 | 674 | 2014 | 2015 | 2537 | 0 | 2362 | 1995 | 2076 | 2405 | 2008 | 2283 | 2017 | 2041 | 52 | 3761 | 2192 | 3377 | 2019 | 3001 | 2813 | 3195 | 2035 |
| SAMEA7578064 | Poultry | Germany | 2015 | 3111 | 3189 | 1138 | 3107 | 3113 | 2373 | 1144 | 1143 | 2620 | 2362 | 0 | 1125 | 2679 | 2856 | 1136 | 2761 | 1147 | 1171 | 2360 | 4165 | 1273 | 3177 | 1147 | 2388 | 3221 | 3773 | 1165 |
| SAMEA7578177 | Poultry | Denmark | 2015 | 2685 | 2763 | 39 | 2681 | 2687 | 1818 | 95 | 94 | 1982 | 1995 | 1125 | 0 | 2220 | 2321 | 87 | 2289 | 48 | 122 | 1993 | 3506 | 558 | 2580 | 88 | 2841 | 2566 | 3104 | 66 |
| SAMEA8227255 | Poultry | Lithuania | 2016 | 2665 | 2767 | 2233 | 2685 | 2693 | 2172 | 2239 | 2238 | 2348 | 2076 | 2679 | 2220 | 0 | 2479 | 2231 | 2289 | 2242 | 2266 | 2074 | 3242 | 2226 | 3351 | 2242 | 3130 | 2295 | 2938 | 2260 |
| SAMEA8227341 | Poultry | Spain | 2016 | 2902 | 3044 | 2334 | 2962 | 2968 | 2479 | 2340 | 2339 | 2126 | 2405 | 2856 | 2321 | 2479 | 0 | 2332 | 2567 | 343 | 2365 | 2403 | 3563 | 2314 | 3494 | 2343 | 2583 | 2680 | 3221 | 2361 |
| SAMEA12943667 | Poultry | Canada | 2015 | 2698 | 2776 | 100 | 2694 | 2700 | 1831 | 64 | 63 | 1953 | 2008 | 1136 | 87 | 2231 | 2332 | 0 | 2300 | 109 | 105 | 2006 | 3521 | 569 | 2595 | 109 | 2852 | 2577 | 3119 | 127 |
| SAMN11444790 | Poultry | United Kingdom | 2017 | 2628 | 2770 | 2302 | 2688 | 2694 | 2345 | 2306 | 2307 | 2353 | 2283 | 2761 | 2289 | 2289 | 2567 | 2300 | 0 | 2311 | 2335 | 2281 | 3798 | 2323 | 2733 | 2311 | 2834 | 3023 | 3116 | 2329 |
| 236996_C2Cspec | Chicken meat | United Kingdom | 2022 | 0 | 397 | 2698 | 329 | 345 | 2471 | 2704 | 2705 | 2843 | 2426 | 3111 | 2685 | 2665 | 2902 | 2698 | 2628 | 2707 | 2733 | 2424 | 3360 | 2858 | 3870 | 2709 | 3131 | 2426 | 3377 | 2725 |
| 245209_C5Cspec | Chicken meat | United Kingdom | 2022 | 397 | 0 | 2776 | 154 | 170 | 2555 | 2782 | 2783 | 2908 | 2510 | 3189 | 2763 | 2767 | 3044 | 2776 | 2770 | 2785 | 2811 | 2508 | 3466 | 2936 | 4006 | 2787 | 3199 | 2523 | 3415 | 2803 |
| 245223_C4Camox | Chicken meat | United Kingdom | 2022 | 2698 | 2776 | 0 | 2694 | 2700 | 1831 | 108 | 107 | 1995 | 2008 | 1138 | 39 | 2233 | 2334 | 100 | 2302 | 43 | 135 | 2006 | 3519 | 571 | 2593 | 101 | 2854 | 2579 | 3117 | 61 |
| 265837_C3CT3 | Chicken meat | United Kingdom | 2022 | 329 | 154 | 2694 | 0 | 88 | 2473 | 2700 | 2701 | 2826 | 2428 | 3107 | 2681 | 2685 | 2962 | 2694 | 2688 | 2703 | 2729 | 2426 | 3384 | 2854 | 3924 | 2705 | 3117 | 2441 | 3333 | 2721 |
| 265839_C4AT1 | Chicken meat | United Kingdom | 2022 | 345 | 170 | 2700 | 88 | 0 | 2479 | 2706 | 2707 | 2832 | 2432 | 3113 | 2687 | 2693 | 2968 | 2700 | 2694 | 2709 | 2735 | 2430 | 3390 | 2860 | 3930 | 2711 | 3123 | 2447 | 3339 | 2727 |
| 265840_C4AT2 | Chicken meat | United Kingdom | 2022 | 2471 | 2555 | 1831 | 2473 | 2479 | 0 | 1837 | 1838 | 2432 | 674 | 2373 | 1818 | 2172 | 2479 | 1831 | 2345 | 1840 | 1864 | 674 | 3379 | 2189 | 3442 | 1842 | 2962 | 2435 | 3091 | 1858 |
